# Predictive value of brain 18F-FDG PET neuroimaging in a mouse model of sudden death

**DOI:** 10.64898/2026.09.14.751341

**Authors:** Elena Marcos-Macías, Juan Miguel Godoy-Corchuelo, Rubén Fernández de la Rosa, Michelle Estefanía Guamán-Chulunchana, Arantxa García-Martín, Luciano Fonseca Lemos de Oliveira, Rafael Dias de Brito Oliveira, Enrico de Francisco Magnani, Miguel Ángel Pozo, Luis García-García, Mirjam Brackhan, Silvia Corrochano, Pablo Bascuñana

## Abstract

Sudden unexpected death (SUD) is a major cause of mortality in neurological disorders, with poorly understood mechanisms and no reliable biomarkers. Brain–heart axis dysfunction has been implicated in SUD pathogenesis. Using the FUSΔ14 mouse model, we tested whether cerebral glucometabolism, assessed by ¹⁸F-FDG PET, reflects autonomic dysregulation and predicts SUD vulnerability.

¹⁸F-FDG PET/CT and ECG were acquired at 10 weeks of age (FUS^Δ14/Δ14^ n=12, FUS^+/Δ14^ n=14, FUS^+/+^ n=12). Heart rate variability was analyzed in time and frequency domains, brain uptake via volume-of-interest and voxel-wise approaches, and correlations assessed by Pearson’s analysis. A classification-tree model tested SUD predictability within the FUSΔ14/Δ14 group.

Female FUSΔ14/Δ14 mice showed reduced ¹⁸F-FDG uptake in basal forebrain septum, left amygdala, and brainstem, with increased uptake in hypothalamus and right midbrain, alongside cardiac size, metabolic and heart rate variability alterations. 50% experienced SUD; in this subgroup, PET showed cortical hypometabolism (insular, entorhinal) and hypermetabolism in hypothalamus, amygdala, septum, and brainstem. Although brainstem uptake was able to singlehandedly predict SUD in this particular FUS^Δ14/Δ14^ group (AUC=0.787±0.06), cross-validation also highlighted cortical metabolism as a relevant feature (UsageRate=0.38 and 0.44, respectively).

¹⁸F-FDG PET identifies region-specific metabolic signatures of brain–heart axis dysfunction linked to SUD, supporting its potential as a non-invasive biomarker for risk stratification.

**Significance statement:** Sudden unexpected death (SUD) is common in neurological disease, yet there are no reliable tools to identify patients at risk. Using a mouse model that mimics seizures and high mortality rate seen in sudden death conditions such as SUDEP, we show that brain-heart axis glucose metabolism, measured by PET imaging, is altered, revealing disturbances in the brain circuits that control heart rhythm. Animals that died unexpectedly displayed a distinct metabolic signature—reduced cortical activity paired with heightened activity in brainstem and structures involved in autonomic control. Exploratory analyses point to brainstem and cortical metabolism as important predictors of SUD risk. These findings position ¹⁸F-FDG PET as a candidate biomarker of brain–heart axis dysfunction, offering a path toward identifying vulnerable individuals.

## Introduction

Sudden unexpected death (SUD) is a term referred to any, non-traumatic, non-drowning, witnessed or unwitnessed, death occurring rapidly after symptom onset, with no cause of death identified after autopsy [1, 2]. From 2010 to 2020 there was an estimate of 2,583,559 attributed sudden deaths in Europe, corresponding to a 31% increase in age-adjusted mortality in this period [3]. Neurological disorders such as stroke, subarachnoid haemorrhage (SAH), epilepsy, Parkinson’s disease (PD), multiple sclerosis (MS), or amyotrophic lateral sclerosis (ALS) among others, present higher SUD mortality rates compared to the general population [1, 4]. One of the most common neurological-related SUD (neuroSUD) situations is sudden unexpected death in epilepsy (SUDEP), which affects 1-2 per 1000 epilepsy patients a year [5, 6] and is second only to stroke in terms of years of life lost [7]. SUDEP is especially significant since it occurs in rather young individuals, the main risk factor being the presence of non-controlled tonic-clonic seizures [6, 8]. Despite being a relatively common condition, the mechanisms by which neurological disorders, and particularly epilepsy, lead to SUD are poorly understood.

The high comorbidity between brain damage and heart dysfunction has evoked a growing interest in the study of the functional relation between brain and heart —the brain–heart axis—to help elucidate the pathophysiology of neurological disorders and SUD. The main pathway connecting both organs is the autonomic nervous system (ANS), which encompasses peripheral structures, brain stem centers and cortical regions, the latter known as the central autonomic network (CAN). One example is that, in stroke, cardiac consequences are frequent and have been linked to CAN areas, such as the insular cortex [9]. Another example is the cardiac autonomic disfunction in Parkinson’s disease and its likely association with SUDPAR [10]. In this regard, epilepsy patients, and particularly high SUDEP-risk individuals, are well known to exhibit altered sympathetic signaling to the heart as well as heart-rate variability abnormalities related to both sympathetic and parasympathetic dysregulation [5, 7]. Experimental evidence has further demonstrated the association between alterations in CAN regions and cardiac dysfunction. For example, seizure-simulating electrical stimulation of the insular cortex (IC) produces autonomic symptoms as well as apnea and arrhythmias [11]. In a preclinical stroke study, researchers linked neuroinflammation in the IC to impaired cardiac function [12]. Moreover, the lithium–pilocarpine rat model of epilepsy, which exhibits electrocardiographic abnormalities, shows both glucometabolic [13] and neuroinflammatory [14] alterations in CAN regions. Thus, it is plausible that the study of the brain-heart axis and autonomic imbalance might shed light upon the mechanisms leading to neuroSUD.

The FUSDelta14 (FUSΔ14) physiological knock-in mouse model carries a mutation in the fused in sarcoma (FUS) gene [15, 16]. In humans, FUS mutations are usually related to an aggressive juvenile form of ALS [17, 18] but have also been linked with frontotemporal dementia [18] and seizures [19]. Homozygous FUSΔ14 mice show smaller brains with altered morphology, reduced neuronal numbers and increased gliosis [15]. Interestingly, homozygous mice present spontaneous seizures related to high rates of SUD —up to 40% in females between 10 and 16 weeks of age—before the onset of motor symptoms [15].

One of the main hurdles in neuroSUD research, and in SUDEP as its most frequent form, is the lack of reliable biomarkers to assess death-risk [20]. ^18^F-fluorodeoxyglucose positron emission tomography (^18^F-FDG PET) is widely used in neurological disorders for diagnostic purposes, enabling early detection of functional changes and multiorganic imaging. This makes the technique particularly suitable for studying the role of the brain-heart axis in SUDEP and for identifying an early biomarker. To date, PET studies in high SUDEP-risk patients, defined by the presence of uncontrolled tonic-clonic seizures, have revealed increased metabolic rates in brain areas related to autonomic control [21] and reduced rates in the medial frontal cortex [22]. Notably, two FUSopathv case reports using ^18^F-FDG PET found hypometabolism in the left frontotemporoparietal cortex and striatum [23], and left frontoinsular cortex respectively [24], linking Fus mutations with glucometabolic alterations in autonomic-related areas.

Here we interrogate the relation between brain pathology and cardiac alterations leading to SUD in the FUSΔ14 mouse model. We employ a multidisciplinary approach combining ¹⁸F-FDG PET-based analysis of glucometabolic alterations in the brain–heart axis with electrocardiographic assessment of cardiac autonomic dysfunction. Additionally, we evaluate the prognostic value of regional brain metabolic patterns in predicting SUD, using a classification-tree model as an exploratory machine-learning approach.

## Methods

### Animals

To generate the model, C57BL/6J females carrying the FUSΔ14 mutation in heterozygosity (FUS^Δ14/+^ C57BL/6J) were crossed with DBA/2J males, also heterozygous for the mutation (FUS^Δ14/+^ DBA/2J). Experiments were conducted exclusively with F1 C57/DBA mice, which were genotyped at weaning by PCR. The crosses yielded homozygous (FUS^Δ14/Δ14^), heterozygous (FUS^Δ14/+^) and wild-type (FUS^+/+^) mice.

Adult mice were housed in groups of up to 5 individuals, in cages on a ventilated rack (Tecniplast, Buguggiate, Italy), under controlled temperature (22 ± 2° C) and light (12h light/dark cycle) conditions. Animals had unlimited access to standard laboratory rodent chow (Safe, France) and water, and were given nesting material along with gnawing bricks (MCViso, Spain) as environmental enrichment. All procedures were approved by the Institutional Ethics Committee of Instituto de Investigación Biomédica del Hospital Clínico San Carlos, Madrid (C. I. 19/018-II, 26-11-2019) and were carried out in accordance with relevant animal research regulations (Directive 2010/63/EU and RD53/2013). Following the ARRIVE guidelines and the 3R concept, every possible measure was taken to reduce the number of animals and their suffering.

### Experimental procedure

Adult female mice were assigned to one of three experimental groups according to their genotype: FUS^Δ14/Δ14^ (n = 12), FUS^Δ14/+^ (n = 14) or FUS^+/+^ (n = 12). The mice were subjected to an ^18^F-FDG PET scan and simultaneous electrocardiographic (ECG) recording at 10 weeks of age, as well as daily monitoring for SUD throughout the experimental period. The humane endpoint was established at 16 weeks of age to avoid the development of early motor symptoms characteristic of this murine model. After euthanasia by cervical dislocation, the brain and heart were collected, snap-frozen and stored at −80°C.

### 18F-FDG PET imaging procedure and analysis

Mice underwent an ^18^F-FDG PET scan at 10 weeks of age to evaluate brain glucose metabolism. Animals were not fasted before PET imaging. Blood glucose levels were measured right before intraperitoneal radiotracer injection (^18^F-FDG, 5-6 MBq in 0.2 ml saline, Curium Pharma, Spain) and did not vary between groups. Following a 45-minute awake period to ensure brain tracer uptake, animals were placed in a small-animal PET/CT scanner (Albira ARSII, Bruker, Spain) under isoflurane anesthesia (2% in oxygen) for 30 minutes. Both a static 20-minute PET and 10-minute CT were performed with the PET-CT field of view including both head and thorax. PET data was reconstructed by applying the maximum likelihood expectation maximization algorithm, scatter, random and decay corrections whereas the filtered back projection algorithm was used for the CT.

Taking into account that the FUSDelta14 mutation alters brain morphology [15], brain PET images were coregistered to both a standard magnetic resonance imaging (MRI) template (Mirrione) [25] and a custom-made genotype-specific MRI template (see supplementary materials). Using the PMOD 4.1 software (PMOD Technologies, Switzerland), CT images were first coregistered to the MRI templates and then the resulting mathematical transformation was applied to the corresponding SUV-corrected PET images. Then, pre-defined volume of interest (VOI) atlases adapted to each template were applied and ^18^F-FDG uptake ratios to whole brain (SUVR_WB_) calculated. Additionally, voxel-wise comparisons were performed to identify subregional differences. Parametric images of group t-test comparisons were obtained by statistical parametric comparisons (SPM12, University College London). Significance threshold was defined at 0.05 with no correction by multiple comparisons and minimum cluster size was set at 20 voxels.

For cardiac analysis, a VOI was manually created using the radiotracer’s signal intensity drop as the heart’s limit and the CT for anatomical reference for cardiac analysis. SUV correction was also applied.

### ECG procedure and analysis

During the PET-CT procedure, a single-lead ECG recording was acquired using a 3-electrode limb lead configuration (BIOPAC MP160, BIOPAC Systems Inc., United States). The recording was continuously acquired for at least 20 minutes at a sampling rate of 1,000 Hz (AcqKnowledge v5.0 Data Acquisition Software, Biopac Systems, Inc., United States).

Recordings were analyzed offline using the ECG Analysis module of LabChart 7 Pro (ADInstruments, Bella Vista, Australia). Stable, artifact-free sinus rhythm epochs were selected for analysis, whereas ectopic beats, segments with baseline drift, and recordings contaminated by movement or electrical noise were excluded. RR interval series were subsequently exported for time- and frequency-domain heart rate variability (HRV) analysis using CardioSeries v2.7 software, following the methods described by Fanzan et al. [26]. Heart rate (HR), standard deviation of the normal-to-normal RR interval (SDNN), root mean square of successive NN intervals (RMSSD), low-(LF: 0.1–1 Hz) and high-frequency (HF: 1–5 Hz) spectral power, and the LF/HF ratio were calculated.

The QT interval was measured from the onset of the QRS complex to the end of the T wave on signal-averaged complexes, applying murine-specific criteria [27, 28]: in complexes with a positive T wave, the end of the T wave was defined as the point at which the waveform returned to the isoelectric baseline; in complexes with a negative T wave, the end was defined at the nadir of the negative deflection; in complexes showing J–T fusion or biphasic terminal repolarization, the QT interval was extended to the terminal end of the final T-wave component. The corrected QT interval (QTc) was calculated using the Mitchell formula: QTc = QT / √(RR/100), where QT and RR are expressed in milliseconds [29]. Given the recognized limitations of heart rate correction in rodents, uncorrected QT and RR values were also retained for analysis and reported alongside QTc [30].

### [3H]PK11195 autoradiography procedure and analysis

14 µm brain (coronal; bregma +0.86, −1.70, −2.80 mm) and heart (transversal) sections, were obtained at −20° using a cryostat (Leica CM1850, Leica Biosystems, Nubloch, Germany), mounted onto Superfrost Plus slides (Thermo Scientific, Dreieich, Germany) and stored at −80°C. Tissue inflammation was assessed by [^3^H]PK11195 (Perkin Elmer, Rodgau, Germany) autoradiography to evaluate translocator protein (TSPO) expression. Sections were dried on a hot plate at 37°C for approximately 5 min, preincubated at room temperature (RT) in Tris-HCl (50 mM, pH 7.4) for 15 minutes, and subsequently incubated at RT in [3H]PK11195 1 nM for 60 min. After two 5-minute washes in ice-cold preincubation buffer, followed by a brief dip in ice-cold distilled water, slides were air-dried and exposed to radiographic film (Western blotting films, Electron Microscopy Sciences, Morgantown, USA) for 6 weeks [31]. Developed films were placed on a light box (Kaiser Prolite 5000, Kaiser Fototechnik, Buchen, Germany) and images were obtained using a Leica DFC425 camera attached to a Leica MZ6 magnifying glass. Images were analyzed using PMOD 4.1 (PMOD Technologies, Switzerland) and ImageJ software, obtaining optical density values.

### Binary classification tree

Aiming to identify biomarkers of SUD risk and given the limited predictive power of linear regression, we implemented a binary classification tree model. This supervised statistical learning approach enables the capture of non-linear relationships within the dataset while remaining highly interpretable [32]. In the training dataset, which included only FUS^Δ14/Δ14^ mice, SUD status was used as the dependent variable and regional ^18^F-FDG uptake values as predictors. The split criterion of the decision tree was Gini’s diversity index, a measure of dataset impurity, defined as:

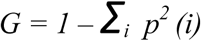

where *p*(*i*)is the relative frequency of the *i*th class within the dataset [33]. In this context, impurity reflects the distribution of class labels, with maximum impurity corresponding to a uniform distribution of labels and minimum impurity corresponding to cases in which all observations belong to the same class. Accordingly, the decision tree selects the splitting variable and threshold at each node that minimizes impurity in the resulting child subsets.

Model performance was evaluated using 4-fold cross-validation repeated over 1000 iterations. Error rates (ER) and area under the curve (AUC) were computed for each iteration, but only best, worst, and closest-to-median performing iterations are reported. The 95% confidence interval for the AUC was derived from the 2.5th and 97.5th percentiles of the AUC distribution across all valid runs. Importance for each predictor was computed as the sum of the impurity reduction in each node where the predictor was used as a splitting variable:

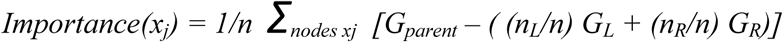

where *x_j_*is the predictor, *nodes x_j_* the number of nodes where the x_j_ predictor is used, *n* the total number of observations in the dataset, *n_L_* and *n_R_* are the number of samples going left and right respectively, *G_parent_* is the Gini index for the parent node and *G_L_* and *G_R_*are the Gini indexes for the left and right child nodes [33]. All analysis was done using MATLAB R2024a (MathWorks Inc., USA). An open-source implementation is provided to ensure full reproducibility (https://github.com/elena-marcos-macias/classificationTree_FusHOM.git).

### Statistics

All data are presented as mean ± standard deviation, unless stated otherwise. Statistical analysis was performed using Prism 10 (GraphPad Software, USA) or MATLAB R2024a (MathWorks Inc., USA). Normality was assessed using the Shapiro-Wilk test. To identify differences between experimental groups, one-way ANOVA, unpaired Student’s t-test or their non-parametric counterparts were applied according to the characteristics of each specific comparison. Bonferroni correction was used in *post-hoc* analysis. *P* values under 0.05 were considered statistically significant.

Pearson’s coefficient was used to evaluate possible correlations between cardiac autonomic function measurements and ^18^F-FDG uptake. Linear regression relations were also explored using a statistical parametric mapping approach. Statistical significance was considered at p < 0.05.

## Results

### FUSΔ14 mutation alters metabolism in the brain-heart axis

First, we sought to describe whether metabolic activity in the brain-heart axis differed between genotypes. Regarding brain-PET, when coregistered to their genotype-specific MRI template and normalized to whole brain uptake (SUVR_WB_), the three different genotypes show statistically significant differences in the following brain regions related to the CAN: basal forebrain septum (BFS) (F = 7.15; p = 0.003), brain stem (BS) (F = 9.132; p = 0.0007), hypothalamus (HYP) (F = 4.61; p = 0.017), left amygdala (l-AMY) (F = 4.97; p = 0.013) and right midbrain (r-MB) (F = 4.27; p = 0.022). The variance analysis also showed differences in other non-autonomic-related areas: central gray (F = 5.77; p = 0.007), superior coliculi (F = 5.63; p = 0.008), and inferior coliculi (right: F = 11.47; p = 0.0001; left: F = 4.86; p = 0.014) (Fig. 1C).

**Fig. 1.**
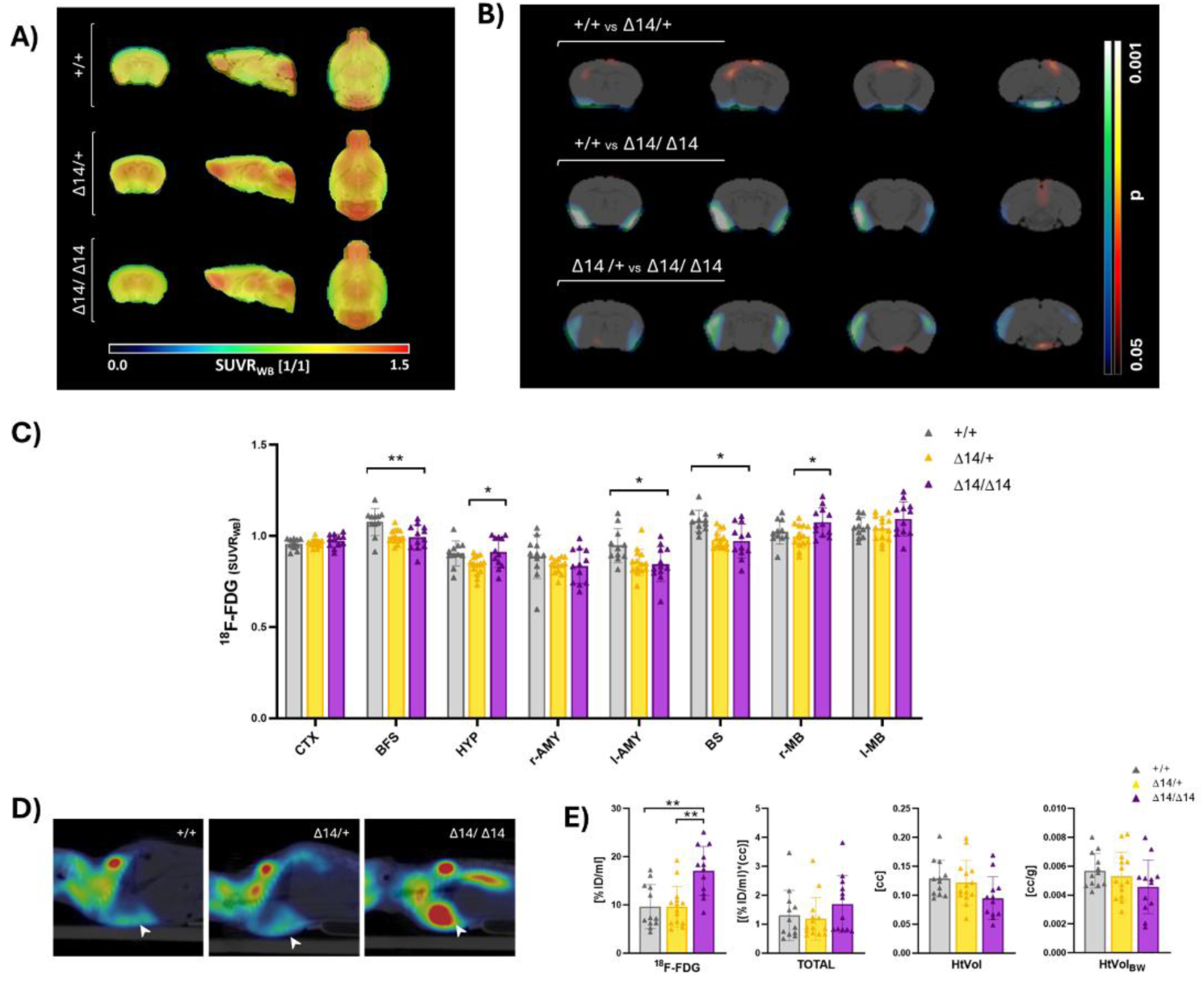
PET imaging analysis of brain and heart ^18^F-FDG uptake. **(A)** Averaged SUVR_WB_ brain PET images per genotype. **(B)** Parametrical images (right hemisphere to the right) reveal bilateral amygdalocortical hypometabolic clusters in FUS^Δ14/Δ14^ when compared to the other two genotypes. Significant p values are depicted in blue-green (lower uptake/binding) and hot (higher uptake/binding) scales. **(C)** Bar graph showing normalized uptake in brain ROIs. FUS^Δ14/Δ14^ display lower uptake values in CAN-related areas than their wild type counterparts. **(D)** Representative SUV heart images per genotype, depicting a higher heart uptake in homozygous individuals. **(E)** FUSΔ14/Δ14 exhibit higher heart metabolic rates, along with smaller cardiac volumes. Data is shown as mean ± standard deviation; one-way ANOVA followed by post hoc Bonferroni test; horizontal lines indicate statistical significance in post hoc analysis; * p < 0.05; ** p < 0.01. CTX, cortex; BFS, basal forebrain septum; HYP, hypothalamus; AMY, amygdala; BS, brain stem; MB, midbrain.

*Post-hoc* analysis shows that, within CAN-related regions, FUS^Δ14/Δ14^ mice exhibit reduced ^18^F-FDG uptake in the BFS (0.992±0.065 vs 1.0765±0.071; p = 0.006), l-AMY (0.854±0.091 vs 0.948±0.088; p = 0.023), and BS (0.971±0.0892 vs 1.081±0.0567; p = 0.002) compared to wild-type mice. In these regions, FUS^Δ14/+^ animals display intermediate metabolic values that are closer to those of FUS^Δ14/Δ14^ mice. FUS^Δ14/Δ14^ mice also show increased uptake in the HYP (0.911±0.0794 vs 0.838±0.0502; p = 0.029) and r-MB (1.076±0.0749 vs 0.997±0.06; p = 0.02) relative to heterozygous animals, with wild-type mice exhibiting intermediate values in both regions (Fig. 1A and C).

Statistical parametric mapping (SPM) analysis not only corroborates this spatial pattern but also reveals additional differences that are not confined to predefined regions. FUS^Δ14/Δ14^ animals, when compared to the other two groups, display two clusters of decreased ^18^F-FDG uptake that start from the right and left AMY respectively and continue dorsally including the piriform (PIRC) and IC. In addition, FUS^Δ14/+^ mice show a cluster of voxels with greater uptake in the supra-hippocampal region relative to FUS^+/+^ mice (Fig. 1B).

For the heart VOI, FUS^Δ14/Δ14^ mice showed statistically significant differences (ANOVA: F = 10.677, p = 0.0002) with higher metabolic rates when compared with FUS^+/+^ (p = 0.002) and FUS^Δ14/+^ individuals (p = 0.0005). However, when uptake was normalized by volume, variance analysis did not reach the threshold for statistical significance. Homozygous animals also displayed a trend toward reduced heart volumes (HtVol), although this effect was attenuated following normalization to body weight (HtVol_BW_) (Fig. 1E).

### 18F-FDG brain PET identifies SUD-suffering mice in high SUD-risk and general populations

Consistent with previous research, homozygous individuals exhibited a 50% mortality rate, while in heterozygous and wild-type mice it remained below 10% at the experimental endpoint. Once metabolic differences between genotypes were characterized, we then aimed to discern between mice that suffered spontaneous deaths during the course of the experiment and those that survived within the high SUD-risk population, i.e. FUS^Δ14/Δ14^ females.

Amongst FUS^Δ14/Δ14^ animals, those that died (SUD^Δ14/Δ14^) showed significantly increased metabolism in BFS (1.03±0.05 vs 0.954±0.06; p = 0.046), HYP (0.971±0.06 vs 0.851±0.06; p = 0.0015), r-AMY (0.897±0.06 vs 0.771±0.08; p = 0.012) and BS (1.049±0.05 vs 0.894±0.05; p = 0.0002) compared to surviving animals. In contrast, they display lower metabolic rates in regions such as CTX (0.946±0.03 vs 1.001±0.01; p = 0.0015) and l-InfCOL (1.024±0.04 vs 1.102±0.04; p = 0.006) (Fig. 2A). The SPM analysis confirmed this spatial distribution showing a dorsal decreased-uptake region with slight lateralization to the left in SUD^Δ14/Δ14^ animals. They also displayed a ventral area with right lateralization, which includes the r-AMY, r-PIRC and r-IC, with higher metabolic rates (Fig. 2B).

**Fig. 2.**
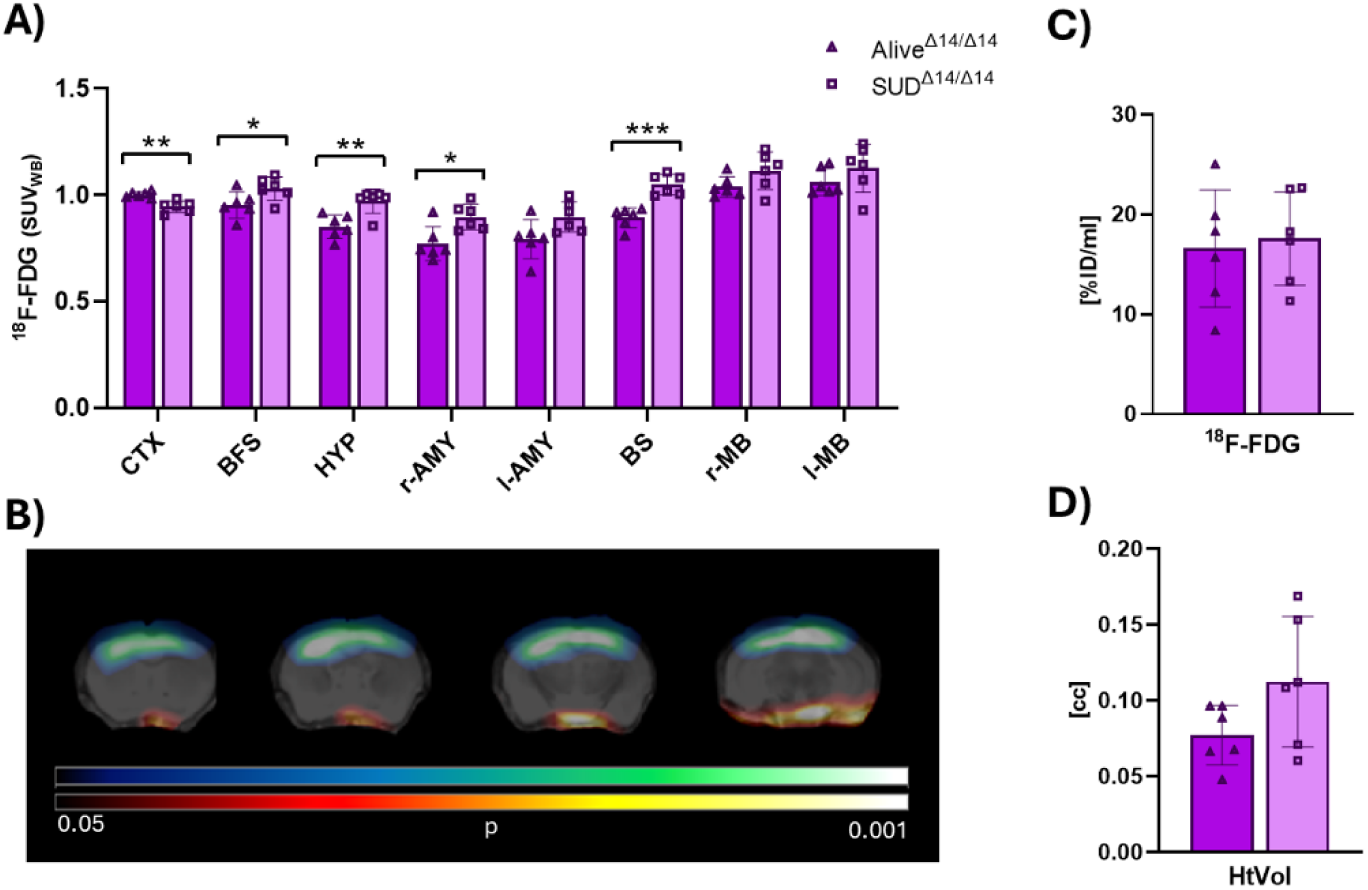
Imaging analysis of brain and heart ¹⁸F-FDG PET comparing SUD and surviving mice. **(A)** SUD^Δ14/Δ14^ mice exhibit higher metabolic rates in CAN-related areas and lower metabolic activity in the cortex compared to surviving homozygous animals. **(B)** Parametric images (right hemisphere displayed on the right) reveal ventral hypermetabolic and dorsocortical hypometabolic clusters that distinguish SUD^Δ14/Δ14^ from Alive^Δ14/Δ14^ mice. Significant p values are depicted in blue-green (lower uptake/binding) and hot (higher uptake/binding). (C–D) Cardiac PET analysis shows no differences in either FDG uptake or cardiac volumes between the two groups. Data are presented as mean ± standard deviation; statistical comparisons were performed using t-test analysis; horizontal lines indicate statistical significance; *p < 0.05; **p < 0.01. CTX, cortex; BFS, basal forebrain septum; HYP, hypothalamus; AMY, amygdala; BS, brainstem; MB, midbrain.

No differences were found between SUD^Δ14/Δ14^ and Alive^Δ14/Δ14^, nor SUD and Alive in general, regarding the heart’s uptake or volume (Fig. 2C and D).

### ECG measures do not distinguish genotype or SUD outcome

Variance analysis of ECG measures across the three genotypes yielded no statistically significant differences in any of the assessed parameters, including heart rate, RMSSD, SDNN, LF/HF ratio, and corrected QT interval (QTc) (Fig. 3A). Similarly, none of the selected ECG variables were able to differentiate between mice that suffered SUD and those that survived among the high SUD-risk FUS^Δ14/Δ14^ population (Fig. 3B).

**Fig. 3.**
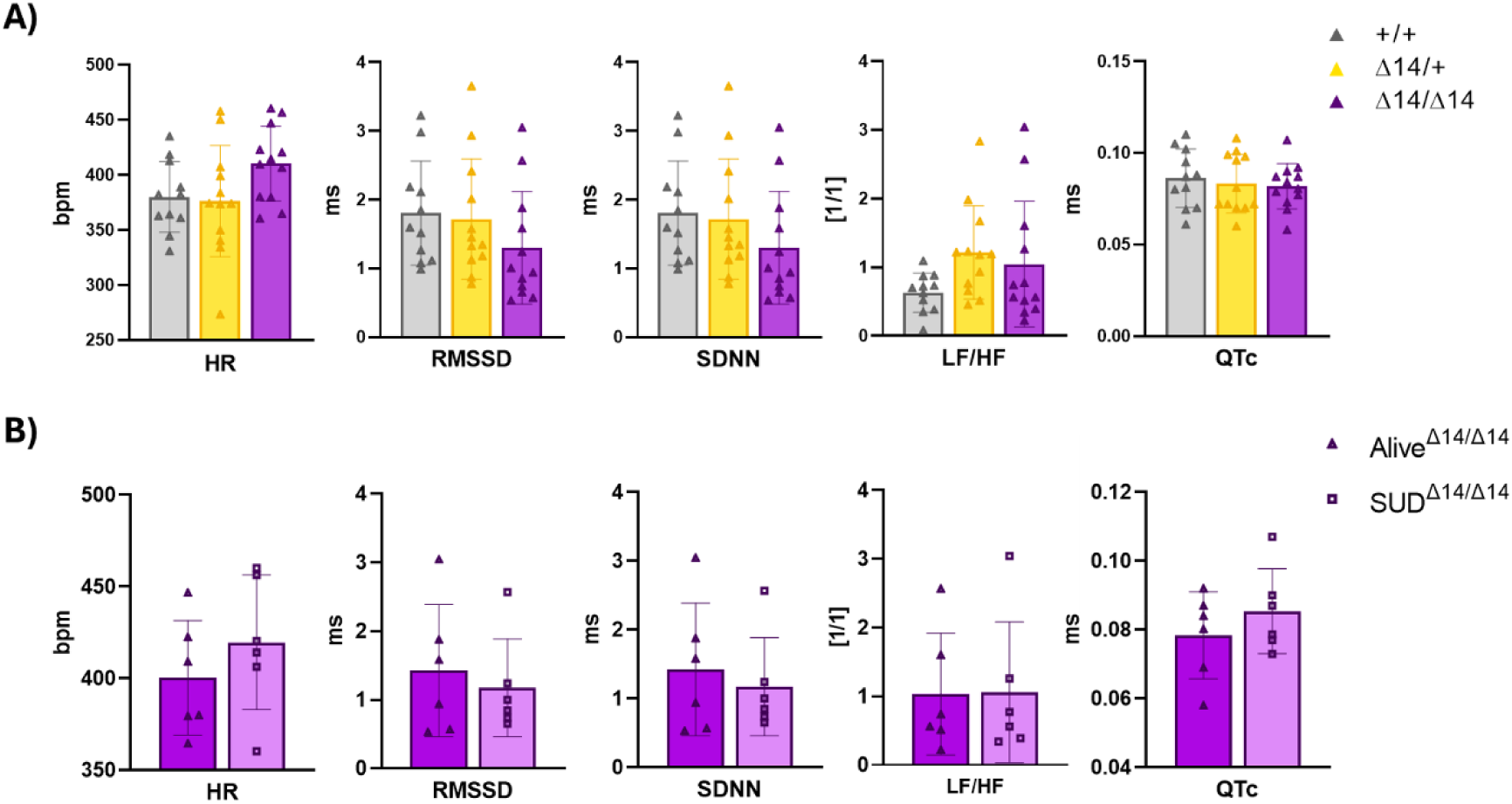
ECG analysis shows no statistically significant differences in the most common HRV parameters. Graphs **(A)** displays comparisons across genotypes (one-way ANOVA) and **(B)** between FUS^Δ14/Δ14^ that suffer SUD and those that survive (t-test). Data are presented as mean ± standard deviation; horizontal lines indicate statistical significance; *p < 0.05. HR, heart rate; RMSSD, root mean squared of successive differences in RR intervals; SDNN, standard deviation of normal-to-normal RR intervals; LF/HF, low frequency/ high frequency ratio; QTc, corrected QT interval; SUD, sudden unexpected death.

### Glucometabolic differences across genotypes occur independently of TSPO-related inflammation

Whenever possible, autoradiographic assessment of TSPO expression —as a surrogate marker of inflammation—was conducted in brain and heart regions exhibiting altered ¹⁸F-FDG uptake. [3H]PK11195 binding was comparable across all different genotypes in the analyzed brain and cardiac regions (Fig. S1).

### FUS^Δ14/Δ14^ brain metabolism correlates best with HR and LF/HF

A linear regression analysis was conducted between ECG parameters and brain ^18^F-FDG uptake. Neither the overall cohort (all mice) nor the high SUD-risk group (FUS^Δ14/Δ14^ mice) showed strong associations between the selected brain regions and ECG metrics. In FUS^Δ14/Δ14^ mice, a mild negative correlation was found between HR and l-STR metabolism (R^2^ = 0.496; p =0.011).

In FUS^Δ14/Δ14^ animals, SPM analysis identified correlations between HR and two brain voxel clusters: a negative correlation with cluster A1, corresponding to a subregion of the left striatum (R² = 0.7187; p = 0.0005), and a positive correlation with cluster A2 located in the BS (R² = 0.3991; p = 0.0275) (Fig. 4A and B). The LF/HF ratio showed positive correlations with several clusters: A3, nearly overlapping the left insular cortex (R² = 0.7359; p = 0.0004) and A4, in the parietal cortex (R² = 0.5377; p = 0.0067) (Fig. 4A and C). However, these correlations did not differentiate SUD^Δ14/Δ14^ and Alive^Δ14/Δ14^ along the linear regression line.

**Fig. 4.**
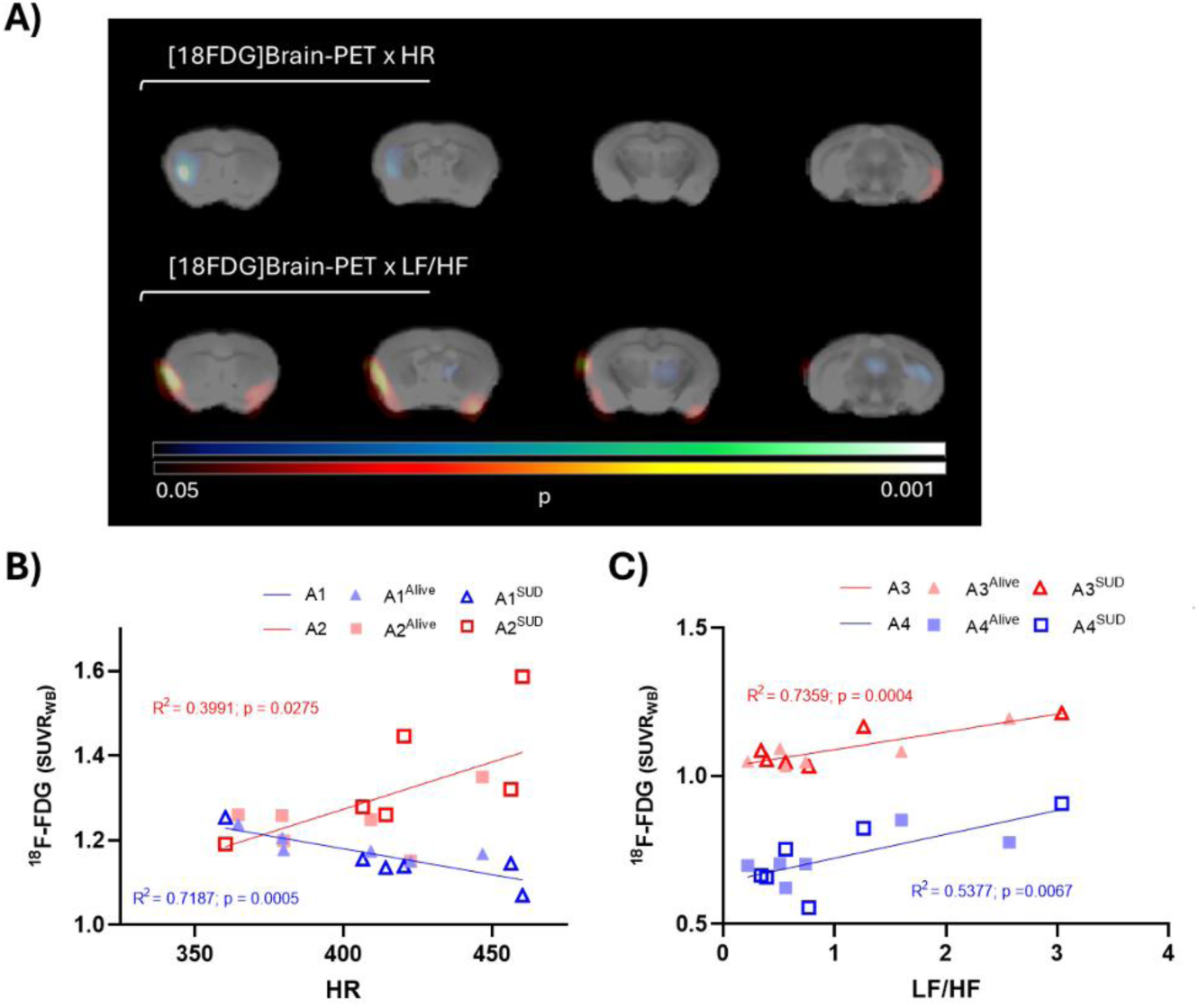
SPM analysis reveals linear correlations between HRV measurements and clusters in the brain PET. **(A)** Statistical parametric images showing positive (hot scale) and negative (cold scale) correlations between brain metabolism and heart rate (top image) and LF/HF (bottom image). **(B)** Strong correlations between brain metabolism and heart rate. A1, subregion of the left striatum; A2, located in the brain stem. **(C)** Positive correlations between brain metabolism and LF/HF. A3, left insular cortex; A4, located in the parietal cortex.

### Classification tree proposes brainstem and cortex ^18^F-FDG uptake as the main predictors of SUD

The classification tree model achieved maximal impurity reduction with a single split node, using brainstem ¹⁸F-FDG uptake as the sole predictor of SUD (BS ≥ 0.966) (Fig. 5A). Cross-validation yielded preliminary performance metrics, with mean error rates of 0.213 ± 0.06 and AUC of 0.787 ± 0.06 (95% CI 0.75 - 0.833) (Table 1). The distribution of said metrics further supports this idea since median values coincide with best performance scenarios (Table 1 and Fig. 5B-D). However, predictor selection was not consistent across all trees generated during the cross-validation process (4-fold × 1000 iterations). Analysis of variable importance across iterations revealed that CTX uptake was the most frequently selected predictor (usage rate = 0.44), despite not being included in the tree derived from the whole dataset (Table 2).

**Fig. 5.**
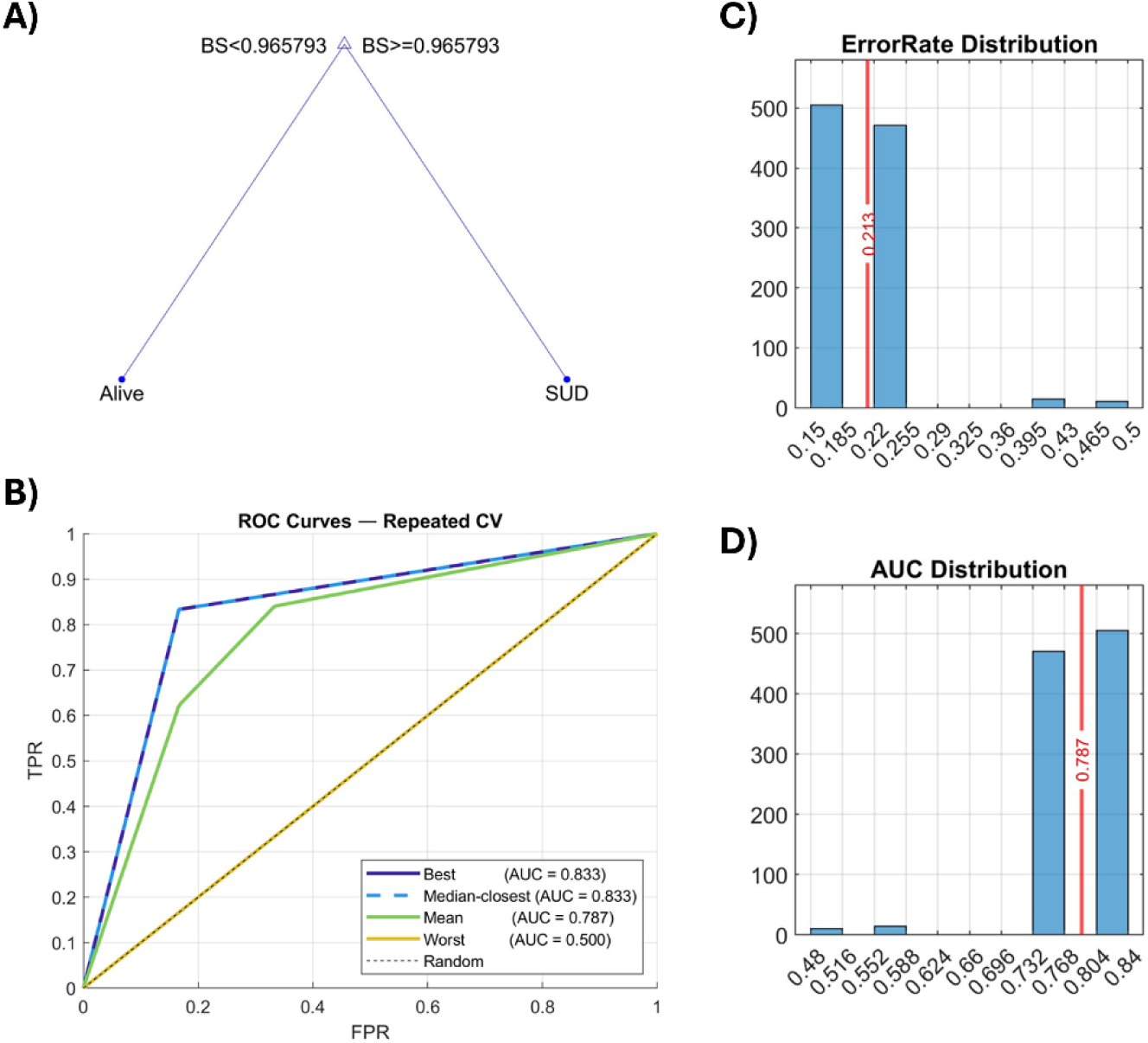
Classification tree: predictors and cross-validation metrics. **(A)** This single node classification tree figure represents the brainstem’s uptake threshold predicting SUD. **(B)** Receiver Operating Characteristic (ROC) curves from best, worst and closest to median performing scenarios from all the possible cross-validation-created trees. Mean ROC curve is separately calculated and does not correspond with a cross-validation scenario. TRP, true positive rate; FPR, false positive rate. **(C and D)** Histograms representing the distribution of all the error rate and AUC values, respectively, obtained from each cross-validation tree. The red vertical line represents the mean value.

**Table 1.**
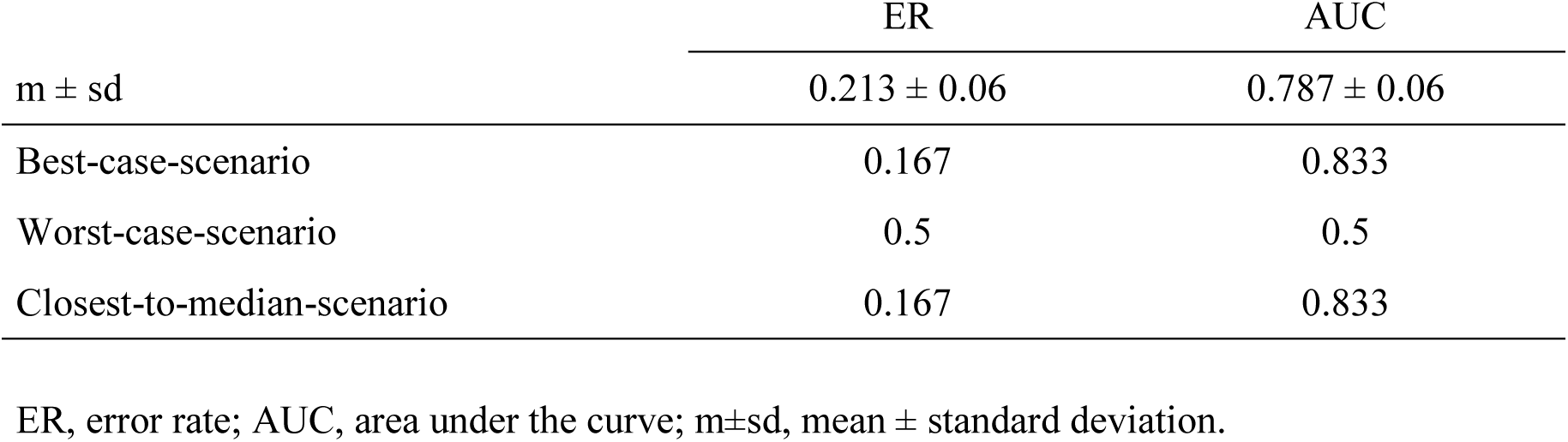
Cross-validation performance.

|  | ER | AUC |
| --- | --- | --- |
| m ± sd | 0.213 ± 0.06 | 0.787 ± 0.06 |
| Best-case-scenario | 0.167 | 0.833 |
| Worst-case-scenario | 0.5 | 0.5 |
| Closest-to-median-scenario | 0.167 | 0.833 |

ER, error rate; AUC, area under the curve; m±sd, mean ± standard deviation.

**Table 2.** Predictor’s Importance throughout cross-validation.

|  | rSTR | lSTR | CTX | rAMY | BS |
| --- | --- | --- | --- | --- | --- |
| Imp (m±sd) | 0.0024 ± 0.034 | 0.003 ± 0.039 | 0.22 ± 0.248 | 0.083 ± 0.19 | 0.192 ± 0.243 |
| Imp (med) | 0 | 0 | 0 | 0 | 0 |
| UsageRate | 0.005 | 0.006 | 0.44 | 0.166 | 0.38 |
| ValuesTaken | [0, 0.5] | [0, 0.5] | [0, 0.5] | [0, 0.5] | [0, 0.5] |

Imp, Importance; m±sd, mean ± standard deviation; med, median; UsageRate, rate of times a predictor was used throughout the cross-validation process; ValuesTaken, possible Importance values the predictor has taken throughout the cross-validation process; rSTR, right striatum; lSTR, left striatum; CTX, cortex; rAMY, right amygdala; BS, brain stem.

## Discussion

Sudden unexpected death is a relatively common outcome in neurological disorders, particularly in epilepsy. The strong comorbidity between brain injury and cardiac dysfunction has steered research in this area toward the brain–heart axis and the role of the autonomic nervous system in neuroSUD and, particularly in SUDEP, pathophysiology. Here, we used a multidisciplinary ^18^F-FDG PET/ECG approach to interrogate brain-heart axis function in the FUSDelta14 murine model, as well as explore the possibility of using brain glucometabolism as biomarker for SUD risk in neurological conditions, using linear and non-linear analysis methods. The present study demonstrates that FUS^Δ14/Δ14^ mice exhibit distinct glucometabolic alterations in autonomic-related brain regions and increased cardiac metabolic rates, which together recapitulate key features of the brain–heart axis dysfunction observed in high-SUDEP-risk patients. Critically, among homozygous animals, those that experienced spontaneous death could be prospectively identified by their brain ¹⁸F-FDG PET profile, particularly brainstem and cortical uptake, while conventional ECG and HRV parameters failed to distinguish SUD cases within a high SUD-risk population—i.e. FUS^Δ14/Δ14^ females.

SUDEP is the most frequent form of neurological-related SUD, currently representing the second leading neurological cause of years of life lost [7], affecting relatively young individuals. Despite its clinical impact, SUDEP—as well as sudden death (SUD) associated with other neurological disorders—still lacks a specific preventive treatment, a limitation that is likely attributable to the absence of reliable biomarkers capable of identifying individuals at risk [20]. At present, clinical studies assessing SUDEP risk are scarce and largely rely on the cumulative assessment of clinical risk factors, most commonly through the SUDEP-7 inventory [22, 34], or merely on the presence and frequency of uncontrolled tonic–clonic seizures [21, 35, 36]. The reliance on such broad and indirect criteria has contributed to inconsistent findings across clinical studies. Nevertheless, a recurrent observation is the involvement of brain regions associated with autonomic regulation, accompanied by evidence of autonomic imbalance, most frequently manifested through ECG alterations [2, 7, 33, 37].

Regarding brain-heart axis functionality in the FUSDelta14 murine model, the present study shows that the FUS^Δ14/Δ14^ mice exhibit regional brain hypometabolism, particularly in the BFS, BS, piriform and insular cortices, and amygdalae, while showing increased ¹⁸F-FDG uptake in the heart. This brain hypometabolic pattern seen in homozygous females partially mimics that of two case reports of patients with FUSopathies who display lower metabolic rates in the left frontotemporoinsular cortex and striatum [23, 24] and a review that identifies anterior frontotemporal hypometabolism in these patients [38]. As previously discussed, studies assessing populations at high SUDEP risk are scarce and rely on vague inclusion criteria, which contributes to inconsistent findings [20], a limitation that also applies to ¹⁸F-FDG brain PET investigations. Within this heterogeneous framework, both hypermetabolic (including the basal ganglia, thalamus, ventral diencephalon, midbrain, pons and deep cerebellar nuclei) [21] and hypometabolic regions (such as the bilateral inferior cortex, including the anterior cingulate; left planum temporale) [21, 22] have been reported in patients at high SUDEP risk. Importantly, all studies converge on the involvement of brain regions responsible for cardiovascular and respiratory regulation. Notably, the regions exhibiting metabolic alterations in the FUS^Δ14/Δ14^ model in this study are also involved in autonomic control [8, 39, 40], thus reproducing key aspects of brain metabolism in high SUDEP-risk patients.

Another key metabolic feature associated with functional Fus protein absence was cardiac hypermetabolism. In a previous study, Ali et.al. described deficient glucose handling and a switch to lipidic metabolic routes in the mice’s muscle and liver tissues [15], suggesting a diversion of glucose resources towards the heart. Here, we found an increase of heart glucose metabolism in a SUD model which warrants further investigation of the metabolic crosstalk between the brain and the heart in SUD.

Surprisingly, despite the cardiac metabolic and morphological alterations observed in homozygous mice, heart rate variability analysis yielded no statistically significant differences in any of the assessed parameters (HR, RMSSD, SDNN, LF/HF ratio, or QTc) across genotypes or between SUD and surviving animals. Although the directional pattern of these measures in FUS^Δ14/Δ14^ animals (higher HR and LF/HF ratio; longer QTc and lower RMSSD values), particularly in SUD-suffering mice, is consistent with an autonomic imbalance profile [41–43], formal statistical support is lacking, likely in part due to the limited sample size within the SUD subgroup. Said electric pattern reproduces that of the “epileptic heart”, an electromechanical dysfunction in the context of chronic epilepsy, also characterized by repolarization abnormalities such as QTc segment elongation [33]. Several studies, including a 2023 metanalysis, revealed that high SUDEP-risk patients also exhibit these HRV modifications. However, similarly to our study, they are unable to successfully identify a cardiac biomarker of SUD risk [37, 41, 44].

When further inspecting the brain and heart interplay in the FUSDelta14 model, we observe that altered regions (IC, AMY and BS) are all part of the CAN, which control sympathetic and parasympathetic autoflow [39]. These areas, particularly the insular cortex, have been linked to autonomic and cardiac disfunction in some of the main neurological disorders related to SUD. In epilepsy, temporoinsular hypometabolism and autonomic manifestations appear hand in hand in both insular [11, 45] and temporal lobe epilepsies [37, 46], the latter being the most prone-to-SUDEP situation among the focal epilepsies [36]. The stroke–heart syndrome, also connects CAN alterations with cardiac dysfunction [36], with preclinical studies further linking insular neuroinflammation to reduced cardiac function [12]. Although direct causality remains to be formally established, these converging findings support a strong relationship between CAN dysfunction and cardiac disruption, which is effectively reproduced in the FUS^Δ14/Δ14^ model, thereby providing a relevant experimental platform to investigate the mechanistic interplay between central autonomic network disruption and cardiac dysregulation in neurological disease. Further supporting this notion, the present study was able to locate a strong positive correlation between left insular metabolism and LF/HF ratio in the FUS^Δ14/Δ14^ population. More specifically, the higher the l-IC metabolic activity, the greater the autonomic imbalance in homozygous mice.

[3H]PK11195 autoradiography, although biased by survival, indicates that these genotype-specific metabolic differences are unlikely to result from alterations in TSPO+ cell populations — predominantly microglia in the brain [47] and proinflammatory cells in peripheral tissues such as the heart [48]. Previous research shows that the FUSΔ14 mutation drives brain progenitor cells towards a glial fate [49], resulting in reduced neuronal numbers and increased astrocytic populations [15]. This shift makes neuronal cell loss a probable explanation for the focal brain metabolic decline seen in FUS^Δ14/Δ14^ mice. In cardiac tissue, higher uptake values in homozygous animals are likely attributable to increased cardiomyocyte activity, as this cell population is thought to account for most ventricular ¹⁸F-FDG uptake [50]. In any case, these results should be interpreted with caution, since tissue from animals that suffered spontaneous deaths is lacking.

In this study, we identified a high SUD-risk group characterized by a 50% mortality rate at 16 weeks of age, consistent with previously reported mortality in this model [15]. This stratification allowed direct comparison between high-risk but surviving individuals and animals that experienced spontaneous death, thereby enabling the exploration of potential predictive biomarkers. Homozygous animals that suffered spontaneous deaths (SUD^Δ14/Δ14^) exhibited a distinct metabolic signature characterized by a ventral hypermetabolic area including BFS, HYP, r-AMY, BS and r-MB, together with a dorsal hypometabolic region covering most of the CTX. In line with these findings, the binary classification-tree model identified BS ¹⁸F-FDG uptake as the sole predictor of SUD. However, given the limited sample size, feature importance derived from cross-validation provides a more robust estimate of predictor relevance [32]. Said analysis indicates cortical uptake as the most important SUD predictor, followed by brainstem metabolism.

Therefore, brain glucometabolism is able to distinguish SUD^Δ14/Δ14^ animals from surviving mice, suggesting that specific metabolic alterations are associated with the lethal phenotype. The only available study in neuroSUD literature analyzing ¹⁸F-FDG PET patterns of actual SUD cases was in the context of SUDEP [22]. Notably, researchers found bilateral medial frontal and temporoparietal hypometabolic foci [22], which does not match the pattern observed in SUD^Δ14/Δ14^ mice. However, this discrepancy may be explained by differences in the comparative framework, as the aforementioned study contrasted actual-SUDEP cases with healthy controls rather than with individuals at high SUDEP risk. Therefore, these differences might be attributable to epilepsy rather than to SUDEP risk. Regarding cardiac analysis, neither heart PET nor the HRV parameters revealed significant differences between SUD^Δ14/Δ14^ and Alive^Δ14/Δ14^ animals. Collectively, these findings suggest that the processes tipping the balance between sudden death and survival are probably located within brain autonomic-related or cortical regions rather than within primary cardiac alterations. Additionally, they help glimpse the potential relevance of brain ¹⁸F-FDG PET imaging as an early biomarker for SUD, as metabolic alterations at ten weeks of age may aid prospective identification of animals that would subsequently experience spontaneous death.

Three main limitations were identified during the course of this study. First, the sample size, particularly in the SUD^Δ14/Δ14^ vs. Alive^Δ14/Δ14^ comparisons, was limited. This constraint stemmed from breeding characteristics of the SUD^Δ14/Δ14^ phenotype, which accounts for approximately 6% of all offspring; increasing the sample size beyond this proportion was therefore not ethically justifiable. Second, ECG recordings were performed under isoflurane anesthesia, which is known to intrinsically affect HR and HRV readings. Moreover, manual adjustment of anesthesia levels throughout the acquisition period may have introduced additional variability in ECG readings. To reduce this limitation, ECG was analyzed using stable artifact-fee segments for the analysis selected by an experienced viewer. All recordings were performed under the same conditions in order to preserve validity of intra-study comparisons. However, with these data we cannot exclude HRV as a potential biomarker of SUD. Finally, because sudden death in these animals is unpredictable, unwitnessed, and predominantly nocturnal, it was not possible to collect brain and heart tissues from SUD cases, preventing direct *in vitro* comparisons between animals that experienced SUD and those at high risk.

### Conclusion

The FUS^Δ14/Δ14^ model reproduces key features of brain–heart axis dysfunction associated with sudden death in neurological disorders, with concurrent cerebral and cardiac metabolic alterations, structural remodeling, and autonomic imbalance. Critically, the metabolic signatures identified — cortical hypometabolism and brainstem hypermetabolism — map onto the central autonomic network and partially mirror findings reported in high-risk SUDEP patients. These results highlight ¹⁸F-FDG PET as a promising non-invasive molecular imaging tool for early SUD risk stratification, with potential applicability in clinical settings where identifying at-risk individuals remains an unmet need.

## Statements & Declarations

### Authors contributions

PB and SC designed and conceptualized the study; EMM, JMGC, RFDR collected the data; EMM, AGM, MEGC, LFLO, RDDBO, EFG and PB analyzed the data; EMM and PB drafted the manuscript. JMGC, LFLO, MAP, LGG, MB and SC contributed to the scientific process, interpretation of the results, and critically revised the manuscript. All the authors approved the final version of this manuscript.

### Ethics approval

All animal experiments were conducted in accordance with the research regulations of the European Union (Directive 2010/63/EU) and Spain (RD53/2013) and with the approval of the local authority (Consejeria de Agricultura, Comunidad de Madrid). All efforts were made to minimize pain, suffering, and the number of animals. The principles outlined in the ARRIVE guidelines and the 3R concept were considered when planning experiments and reporting data.

### Data Availability

Data will be made available from the corresponding author on reasonable request.

## Acknowledgements

This work was funded by grants to PB (Spanish Health Institute Carlos III (PI22/00446) and the “Miguel Servet” program (CP21/00020)) and to SC (Spanish State Research Agency (PID2023-1479-47OB-100)). MB was funded by the ISCIII-HEALTH program from the Spanish Health Institute Carlos III (IHMC22-00032). EMM was founded by the FPU program from the Spanish Ministerio de Ciencia, Innovación y Universidades (FPU24/03044).

## Abbreviations

%ID: % injected dose
¹⁸F-FDG: fluorodeoxyglucose
ALS: amyotrophic lateral sclerosis
AMY: amygdala
ANS: autonomic nervous system
AUC: area under the curve
BFS: basal forebrain septum
BS: brainstem
CAN: central autonomic network
CI: confidence interval
CT: computed tomography
CTX: cortex
ECG: electrocardiogram
ER: error rates
FUS: fused in sarcoma gene
HF: high frequency spectral power
HR: heart rate
HRV: heart rate variability
HtVol: heart volume
HtVol_BW_: heart volume normalized to body weight
HYP: hypothalamus
IC: insular cortex
LF: low frequency spectral power
MB: midbrain
MRI: magnetic resonance imaging
neuroSUD: neurological-related sudden unexpected death
PET: positron emission tomography
PIRC: piriform cortex
QTc: corrected QT interval
RMSSD: root mean square of successive NN intervals
SDNN: standard deviation of the normal-to-normal RR intervals
SPM: statistical parametric mapping
SUD: sudden unexpected death
SUDEP: sudden unexpected death in epilepsy
SUV: standardized uptake value
SUVR_WB_: SUV ratio to whole brain uptake
VOI: volume of interest.

