## Supplementary data for "Predictive value of brain 18F-FDG PET neuroimaging in a mouse model of sudden death"

### Supplementary materials

*Genotype-specific MRI Template construction*

As FUSΔ14 homozygous mice exhibit significant brain atrophy [15], using standard templates to quantify tracer uptake would introduce systematic spatial bias. Therefore, we constructed a custom-made genotype-specific MRI template for FUSΔ14 homozygous mice. For that, we used a total of 20 MRI scans from FUSΔ14 homozygous mice from a previous study [15]. First, we aligned one of the images in all 3 axes. This MRI was used as a reference to coregister the rest of the images. Once aligned, an MRI template was built by averaging all images. The result image was saved in NIfTI format. In addition, for precise ROI quantification, an ROI atlas was created using Mirrione's atlas as reference. Briefly, we normalized the FUSΔ14 template to Mirrione's MRI template in PMOD and saved the inverse transformation. This transformation was applied to Mirrione's ROI atlas. Correct delineation of the regions was visually inspected and, when needed, ROIs were manually modified.

### Supplementary figures

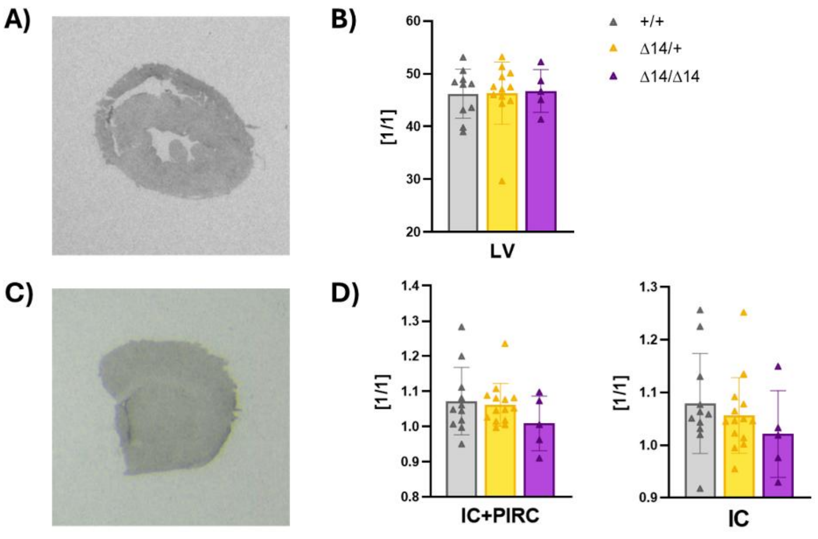


**Fig. S1.** TSPO autoradiography reveals no genotype-dependent differences in brain and heart tissues. **(A)** Representative [$^3$H]PK11195 autoradiography image of cardiac tissue. **(B)** Optical density (O.d.) quantification in the left ventricle (LV). **(C)** Representative [$^3$H]PK11195 autoradiography image of brain tissue. **(D)** Normalized optical density values in the insular cortex (IC) and combined insular and piriform cortices (IC+PIRC). No significant differences were observed across genotypes in any of the analyzed regions.
